# Single cell spatial transcriptomic and translatomic profiling of dopaminergic neurons in health, aging and disease

**DOI:** 10.1101/2023.04.20.537553

**Authors:** Peter Kilfeather, Jia Hui Khoo, Katherina Wagner, Han Liang, Maria-Claudia Caiazza, Yanru An, Xingju Zhang, Xiaoyan Chen, Natalie Connor-Robson, Zhouchun Shang, Richard Wade-Martins

**Author notes:** Joint first authors (Peter Kilfeather, Jia Hui Khoo).

## Abstract

The brain is spatially organized and contains unique cell types, each performing diverse functions, and exhibiting differential susceptibility to neurodegeneration. This is exemplified in Parkinson’s disease with the preferential loss of dopaminergic neurons of the substantia nigra pars compacta. Using a Parkinson’s transgenic model, we conducted a single-cell spatial transcriptomic and dopaminergic neuron translatomic analysis of young and old mouse brains. Through the high resolving capacity of single-cell spatial transcriptomics, we provide a deep characterization of the expression features of dopaminergic neurons and 27 other cell types within their spatial context, identifying markers of healthy and aging cells, spanning Parkinson’s-relevant pathways. We integrate gene enrichment and GWAS data to prioritize putative causative genes for disease investigation, identifying CASR as a novel regulator of dopaminergic calcium handling. These datasets (see: spatialbrain.org) represent the largest public resource for the investigation of spatial gene expression in brain cells in health, aging and disease.

## Introduction

In Parkinson’s disease (PD) there is preferential loss of dopaminergic (DA) neurons of the substantia nigra (SN) pars compacta^1,2^ and intracellular accumulation of ⍺-synuclein. Characteristic motor signs include tremor, bradykinesia, postural instability and rigidity, accompanied by non-motor features, including dementia, depression and psychosis^3^. The disease affects 1% of the global population above the age of 60 years^4^. Age is the biggest risk factor for PD and SN DA neurons may also be lost in healthy aged individuals^5–7^. Overexpression of ⍺-synuclein through locus multiplication causes PD and *in vivo* overexpression of human ⍺-synuclein in the *SNCA-*OVX mouse model recapitulates DA neuron loss^8,9^.

Single-cell RNA-sequencing has advanced our understanding of cell-specific expression in complex tissues, such as brain^10–15^. To isolate individual cells, the tissue is dissociated, resulting in destruction of the tissue architecture and gene expression artifacts^16^. Spatial transcriptomics preserves this architecture, however current sequencing-based spatial technologies do not consistently achieve single-cell or subcellular resolution at high throughput^17^. Fluorescent *in situ* hybridization-based methods offers superior sensitivity at the individual probe level, at lower throughput. Stereo-seq (spatial enhanced resolution omics-sequencing) offers nanoscale resolution spatial expression data, detecting thousands of genes simultaneously^18^.

Single-cell and spatial transcriptomics capture minute quantities of RNA, resulting in lower measurement accuracy compared to bulk RNA-sequencing^19^. Translating ribosome affinity purification (TRAP) enables cell type-specific sequencing with measurement sensitivity comparable to bulk RNA^20–26^. TRAP mRNA is ribosome-bound and engaged in translation, providing a more accurate indicator of protein abundance^27^.

TRAP and RiboTag (a related ribosomal profiling technology) have previously been used to study DA neuron gene expression in mice under healthy conditions and after exposure to the toxin, MPTP^26,28,29^. The dopaminergic neuron aging process has never been characterized using this technology, nor the effects of overexpression of alpha-synuclein.

Single-cell and single-nuclei RNA-sequencing have been used to study DA neuronal expression across a range of species, including humans^11,30,31^ and mice^13,14,32,33^. A comparison of aged and young DA neurons, in a spatial context and complemented with data from other brain cell types, is so far lacking.

In this study, we combined the advantages of Stereo-seq and TRAP to characterize the spatial expression signature of individual cells in mouse brain, with a focus on DA neurons. We optimized a protocol to segment individual cells from Stereo-seq data and subsequently identified 29 cell types across 18 brain sections. By considering the location of each cell, we identified genes with spatially variable expression, such as in SN and ventral tegmental area (VTA) DA neurons. By contrasting DA neuron gene expression with other cell types, we identified strongly specific, yet understudied, markers, including *Slc10a4* and *Cpne7*. Using both long- and short-read sequencing of translating mRNA captured by TRAP, we discovered splice variants specific to DA neurons. We further investigated whether DA axons harbor actively translating ribosomes. To aid the investigation of novel causative genes in PD, we demonstrated how measures of expression specificity from Stereo-seq and TRAP can be used to prioritize candidate genes of interest from GWAS regions. By this process, we identified a novel role for CASR in regulating intracellular calcium handling in DA neurons. We finally compared aged and young brains, revealing a SN-specific loss of DA neurons and expansion of activated microglia with age. Further, we identified a range of age and disease-related expression changes in multiple cell types, including DA neurons, spanning multiple PD-relevant pathways.

## Results

### Integrated transcriptomic profiling in the brain

To combine the advantages of spatial resolution with the sensitivity of the TRAP platform, we generated Rosa26^fsTRAP^ ::DAT^IREScre^ (DAT-TRAP) mice, which express eGFP-L10a in DAT-expressing cells (Figure 1A)^34,35^. DAT-TRAP mice were crossed with *SNCA-*OVX mice and aged to 18 months to investigate the effects of overexpression of human alpha-synuclein and aging on DA neuron gene expression (Figure 1A-B). TRAP samples were prepared from 56 mice, by dissecting the ventral midbrain, dorsal and ventral striatum and incubating each homogenate in paramagnetic beads coated in anti-eGFP antibody (Methods, Figure 1G). Stereo-seq samples were prepared from 18 mice, by cutting 10 µm cryopreserved sections from fresh frozen brains (Methods).

**Figure 1:**
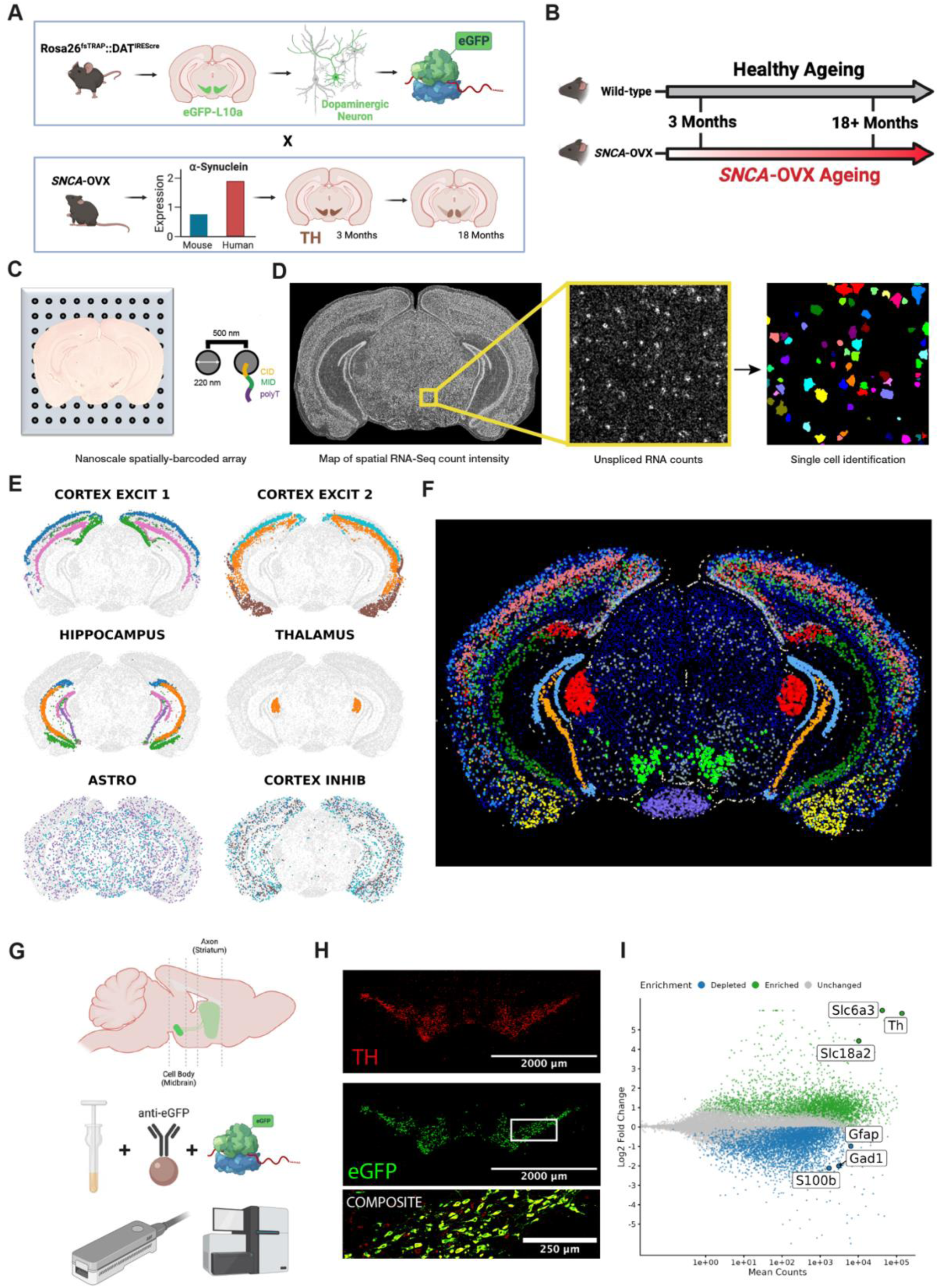
Integration of Stereo-seq and TRAP for high resolution gene expression profiling. A) Transgenic mouse model: Rosa26^fsTRAP^::DAT^IREScre^ mice were crossed with *SNCA*-OVX mice to enable the capture of dopaminergic mRNA in a model of PD. B) Mice were aged to 18+ months in order to study the effect of healthy and Parkinsonian ageing. C) Stereo-seq array, with spatial transcript map from a single brain section. D) Conversion of transcript expression map to segmented individual cells. Segmented cells were subsequently filtered based on transcriptome size and complexity. E-F) Demonstration of spatial compartmentalization of annotated cell types: Spatially distinct populations of the cortex, hippocampus, and thalamus can be visualized. Dispersed cell types were also identified, e.g. astrocytes and inhibitory cortical neurons. A total of 29 distinct cell types were identified. G) Dissection and TRAP processing schematic, before performing short- and long-read RNA sequencing. eGFP-tagged ribosomes in DAT-expressing cells are captured by anti-GFP antibody-coated paramagnetic beads. H) Confirmation of eGFP colocalization with tyrosine hydroxylase (TH), a marker of dopaminergic neurons. I) Demonstration of the specific enrichment of dopaminergic marker genes and depletion of marker genes of other neighboring cell types in DAT-TRAP mRNA (*n* = 56).

**Table 1:**
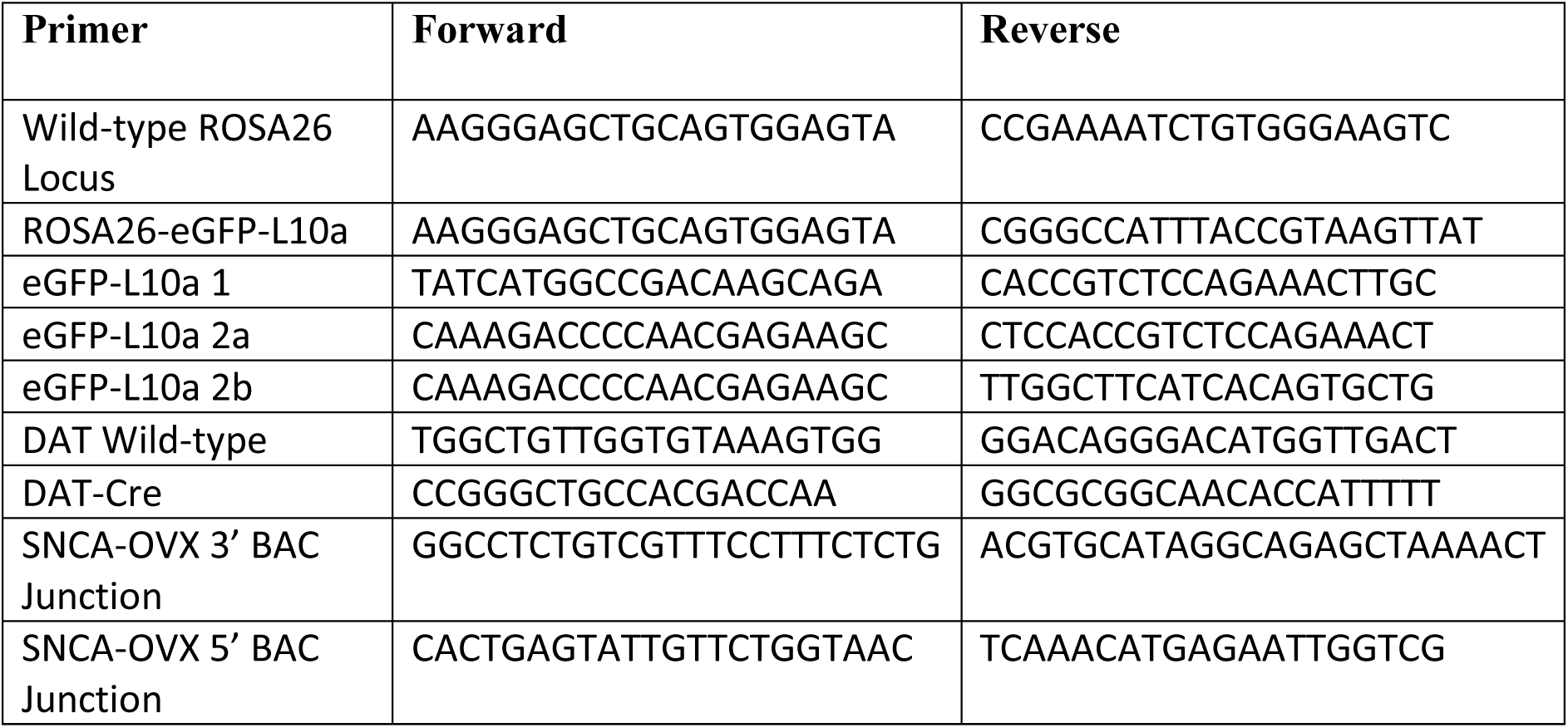
Primer sequences for mouse genotyping.

### Stereo-seq single-cell spatial transcriptomic profiling enables the identification and annotation of distinct cell types in the brain

To generate *in situ* transcriptomic data, 18 cryopreserved mouse brain sections were individually mounted onto DNA nanoball-patterned arrays for library preparation. All sections were analyzed as a single group (encompassing both age groups and genotypes) to validate Stereo-seq analysis methodology (Figure 1C, Methods). We produced a spatial map of transcript detection, to visualize the intensity of RNA capture across each brain (Figure 1D). In each map, we observed distinct anatomical compartments and cell boundaries. We developed a custom image processing pipeline to segment individual cells from Stereo-seq brain sections (Figure 1D, Methods and Supplementary Figure 1 for a detailed example). In the first stage, cells were filtered based on the number of detected genes, to exclude low complexity cells and putative multiplets (Methods, Supplementary Figure 2C). UMAP visualization of all 18 brains confirmed that no obvious batch-related differences were present (Supplementary Figure 2A). We isolated 355,307 high-quality transcriptomes with spatial coordinates from 18 mouse brains. In total, 14,494 genes were detected across all brains, with a median of 626 genes per cell with high-confidence cell segmentation. A summary of the quality control workflow and outcomes is shown in Supplementary Figure 2C.

Spatial transcriptomics enables the identification and annotation of distinct cell types using both expression and location features. Twenty-nine distinct cell types were consistently identified across all brains sections by unsupervised clustering (Methods, Figure 1E-F, summary of all identified cell types in Supplementary Table 1). Cell types were labelled according to both their anatomical localization and the genes most distinctively expressed (marker genes). For example, neurons were distinguished from glia by the expression of *Snap25* and by patterned localization in regions, such as the hippocampus or thalamus (Figure 1E,F). Distinct subpopulations of cells could be visualized, such as neurons of the CA1, CA3, dentate gyrus, and subiculum in the hippocampal region, or GABAergic nuclei within the midbrain. Oligodendrocytes, astrocytes, microglia, and erythrocytes were readily identifiable by marker expression (Oligodendrocytes: *Olig1, Mbp, Sox10, Mog*; Astrocytes: *Gfap, Slc1a3, Atp1a2, Mt3*; Microglia: *Tyrobp/Dap12, Ftl1, Cts(a/b/d/f/h/l/s/z), Aif1, Tmem119, Cd68*; Erythrocytes: *Hba- and Hbb-genes*). Mapping the identity of every cell to its spatial position of origin gives confidence in ascribing greater annotation detail than with expression data alone, e.g. labelling CA1 vs CA3 hippocampal neurons.

### TRAP generates highly sensitive ‘translatomic’ profile of dopaminergic neurons

TRAP complements Stereo-seq by generating a highly sensitive measure of gene expression in a target cell type. In addition, mRNA captured by TRAP is engaged in translation, providing a more accurate readout of gene expression, compared to conventional transcriptomic technologies^27^. For deep characterization of the DA neuron translatome, short- and long-read sequencing were performed on DAT-TRAP samples (Figure 1G). The dissected ventral midbrain and striatum were processed, to enrich for cell body- and putative axon-localized transcripts, respectively. Specific expression of the TRAP transgene was confirmed by immunohistochemical staining for eGFP, which showed distinct colocalization to TH; a marker of DA neurons (Figure 1H). In DAT-TRAP samples, canonical markers of DA neurons were robustly enriched, relative to RNA from bulk tissue homogenate, while markers of other cell types present in the ventral midbrain were depleted (Figure 1I and Supplementary Figure 2D). We compared our DAT-TRAP enrichment data with those of a public DA neuronal RiboTag dataset^28^ and confirmed correlated results, but stronger enrichment of dopaminergic genes using TRAP (Supplementary Figure 2E). Principal component analysis (PCA) of DAT-TRAP samples showed a major distinction between DAT-TRAP samples and tissue homogenate samples, indicating the importance of using cell type-specific RNA over bulk ‘homogenate’ RNA (Supplementary Figure 2F).

### Functionally and spatially distinct populations of dopaminergic neurons are detected by single cell spatial transcriptomic profiling

DA neurons are primarily situated in the ventral midbrain and can be separated into SN and VTA populations by their mediolateral position. SN DA neurons are particularly vulnerable to age-related and Parkinsonian degeneration. By spatially resolving each expression profile, DA neuronal analyses could be focused to those most relevant to PD. We sought to identify DA neurons in our Stereo-seq data and to characterize spatially dependent changes in their expression. In total, 6,378 DA neurons were robustly detected across all 18 brains (Figure 2A). Canonical marker genes, *Th, Slc6a3* (DAT), *Ddc* (Dopa decarboxylase) and *Slc18a2* (VMAT2) were strongly enriched in DA neurons, relative to other cell types (Figure 2B). Highly specific markers were also identified with an underreported role in DA function (e.g. *Slc10a4, Cpne7*): *SLC10A4* is a member of the bile acid transporter family and regulates vesicular uptake of dopamine^36^. *CPNE7* is a calcium-dependent phospholipid binding protein that regulates autophagy and axonal/dendritic extension in other cells^37^. To integrate DA neuron-related data from Stereo-seq and TRAP experiments, a meta-analysis was performed across both enrichment comparisons (Figure 2B). Of the top 100 DA neuron markers identified by Stereo-seq, 98 were also enriched in TRAP samples, confirming the strong concordance of both technologies (Supplementary Figure 3A). The spatial specificity of each marker could be qualitatively confirmed by visualizing the expression of each gene *in situ* (Supplementary Figure 3C).

**Figure 2:**
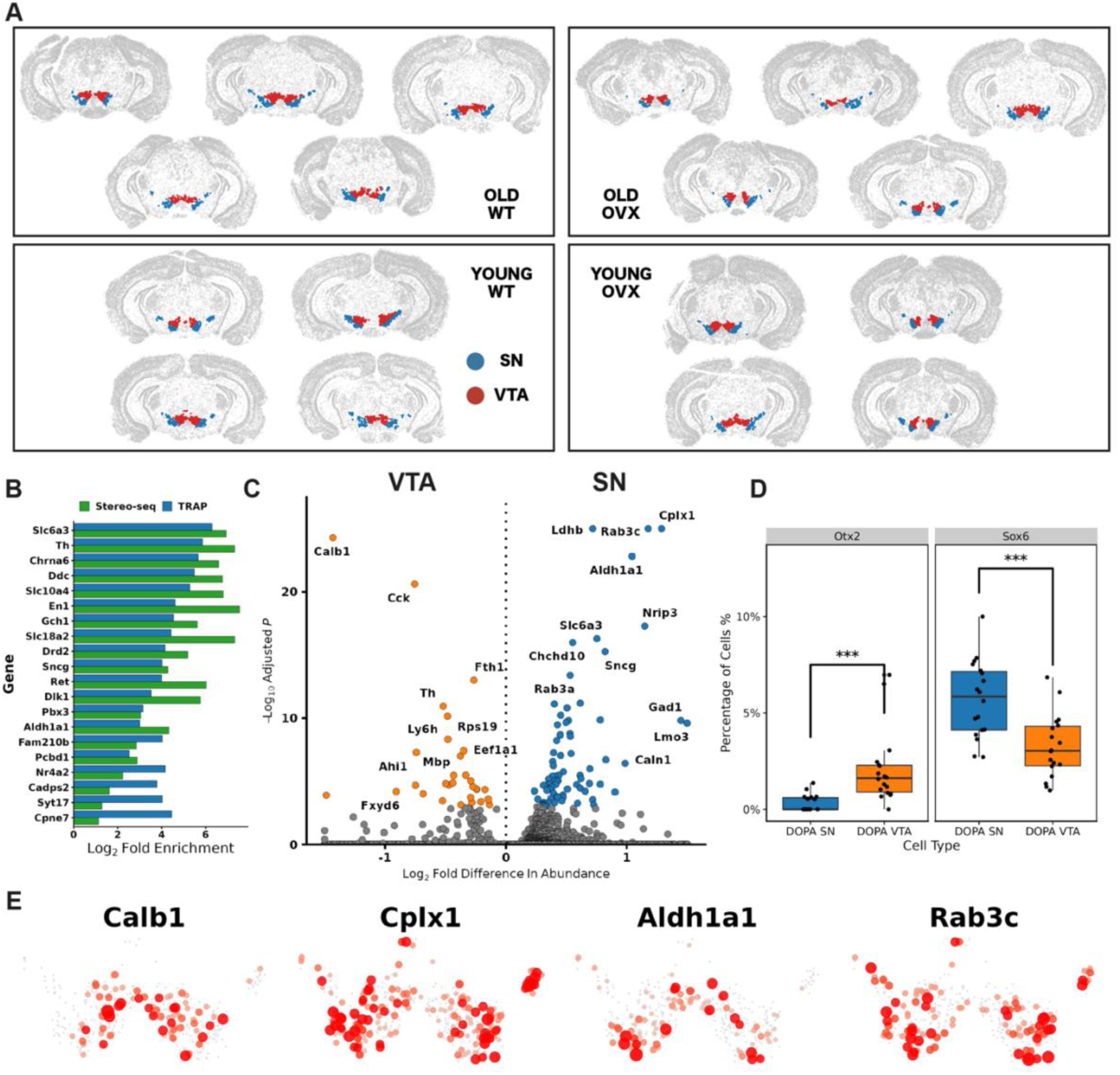
High resolution single cell spatial mapping of sub-populations of DA neurons. **A)** Confirmation of the spatial distribution of cells annotated as dopaminergic neurons. B) Top marker genes of dopaminergic neurons identified from Stereo-seq and *TRAP* data, including less commonly recognized Slc10a4, Gap43, Cpne7. Genes are ranked by the mean fold enrichment in DA neurons from both technologies. C) Markers of SN and VTA neuronal populations. D) Comparison of Otx2 and Sox6 detection rate by region, demonstrating greater expression in VTA and SN, respectively. E) Spatial representation of region-specific marker expression (Calb1 - VTA; Cplx1, Aldh1a1, Rab3c - SN). n = 18 brains, 6378 cells for all Stereo-seq analyses, n = 56 for TRAP analyses.

We used the coordinate positions of Stereo-seq DA neurons to identify 142 genes with significant evidence of spatially variable expression (FDR-adjusted *P* < 0.01, Methods). This revealed a gradient of expression aligning with the SN (e.g. *Cplx1, Nrip3*) and VTA (e.g. *Calb1, Aldh1a1*) of the ventral midbrain (Figure 2C,E). Genes with spatially variable expression in other brain cell types are reported in Supplementary Table 2. 220 differentially expressed genes were identified between the two subpopulations (FDR-adjusted *P* < 0.05), including established (VTA: *Calb1, Calb2*; SN: *Kcnj6* (*Girk2*), *Cplx1*) and putatively novel markers (SN: *Ndnf, Rab3c, Rab6b;* VTA: *Ahi1, Nnat*). Greater *Sox6* and *Otx2* expression was observed in the SN and VTA, respectively, by determining the number of cells in which each was detected (Figure 2D). The spatial visualization of marker gene expression demonstrated that although a SN-VTA distinction is evident, cells from either population can be found in the other brain region. Where possible, the functional study of DA neurons could benefit from stratification of neurons based on the expression of SN-VTA markers.

### TRAP reveals the specificity of transcript expression in dopaminergic neurons

To focus on genes actively translated in DA neurons, a measure of gene enrichment was calculated by comparing the abundance of transcripts in DAT-TRAP mRNA and bulk tissue homogenate mRNA. Across all DAT-TRAP samples, 23,292 genes were detected. Of these genes, 4,828 were found to be more abundant compared to TOTAL RNA. This indicated the subset of the transcriptome that is predominantly translated in DA neurons. We next compared DAT-TRAP samples to published TRAP/RiboTag datasets from glutamatergic and GABAergic neurons, astrocytes, microglia, and oligodendrocytes in ventral midbrain. Overall, 2,504 genes were found to be significantly specifically expressed in DA neurons^22,24,38–40^, (Methods, FDR-adjusted *P <* 0.01) (Supplementary Figure 3B). We leveraged the combination of short- and long-read sequencing technology to profile the specific splice variants that define DA neurons: 1,617 alternatively spliced genes were detected, relative to ventral midbrain RNA (Supplementary Figure 3D,E). Interestingly, splicing was not restricted to genes enriched in DAT-TRAP samples: 817 genes demonstrated evidence of differential transcript usage without gene-level enrichment, suggesting that a substantial component of cell type-specific function could be conferred by splicing and not relative gene-level abundance.

We hypothesized that DA axons locally translate mRNA, due to their extensive projection length into the striatum. We used TRAP to capture putative axonal mRNA from the dorsal and ventral striatum (Figure 1G). In striatal DAT-TRAP samples, we observed an enrichment of 1,803 genes (FDR-adjusted *P* < 0.01), including canonical DA neuron markers, *Th* and *Slc6a3* (DAT) (Supplementary Figure 4A). We compared our enrichment data with a previously reported proteomic characterization of the striatal DA axonal compartment and found significant overlap between enriched genes/proteins (Hypergeometric test, *P* = 1.71e^-29^). The abundance of DA neuron marker genes in striatal DAT-TRAP samples was substantially lower than in midbrain-derived samples, however. We also observed the enrichment of markers of cell types other than DA neurons *(e.g. Gfap, Gad1, Gad2)*. We performed immunohistochemical staining for TH and GFP in mouse brain sections at the level of the striatum (Supplementary Figure 4B). GFP puncta could be identified that colocalized with TH, however the overall signal was sparse.

### Heritability enrichment analysis identifies CASR as a novel regulator of intracellular calcium handling in dopaminergic neurons

Using Stereo-seq and TRAP data, we designed an approach to prioritize candidate genes for sporadic PD investigation. We reasoned that enrichment and specificity measures could be used to partition PD heritability according to causative cell types, as demonstrated previously^33,41^. We measured the cell type-specific enrichment and specificity index (Methods) of genes containing SNPs at an r^2^ > 0.5 and located within ±1 Mb of 107 common risk variants for sporadic PD^42^. 248 out of 303 genes were considered, after retaining genes with mouse homologs (Supplementary Table 3). We observed a broadly neuronal pattern of candidate gene enrichment in Stereo-seq data (Figure 3A), with SN DA neurons demonstrating the greatest mean enrichment. Candidate genes were generally depleted in glial cell types, however *Ctsb, Dpm3, Inpp5f, Rps12, Sbds, Scarb2* and *Stx4a* were commonly enriched between glia and DA neurons (Figure 3A). In TRAP data, DA neurons and oligodendrocytes were jointly found to specifically express the greatest number of candidate genes (Figure 3B). Together, our results indicate a primary role for DA neurons in conferring genetic risk of sporadic PD, however the common enrichment of a minority of genes across distinct cell types also supports cell type-agnostic disease processes, as reported previously^43^.

**Figure 3:**
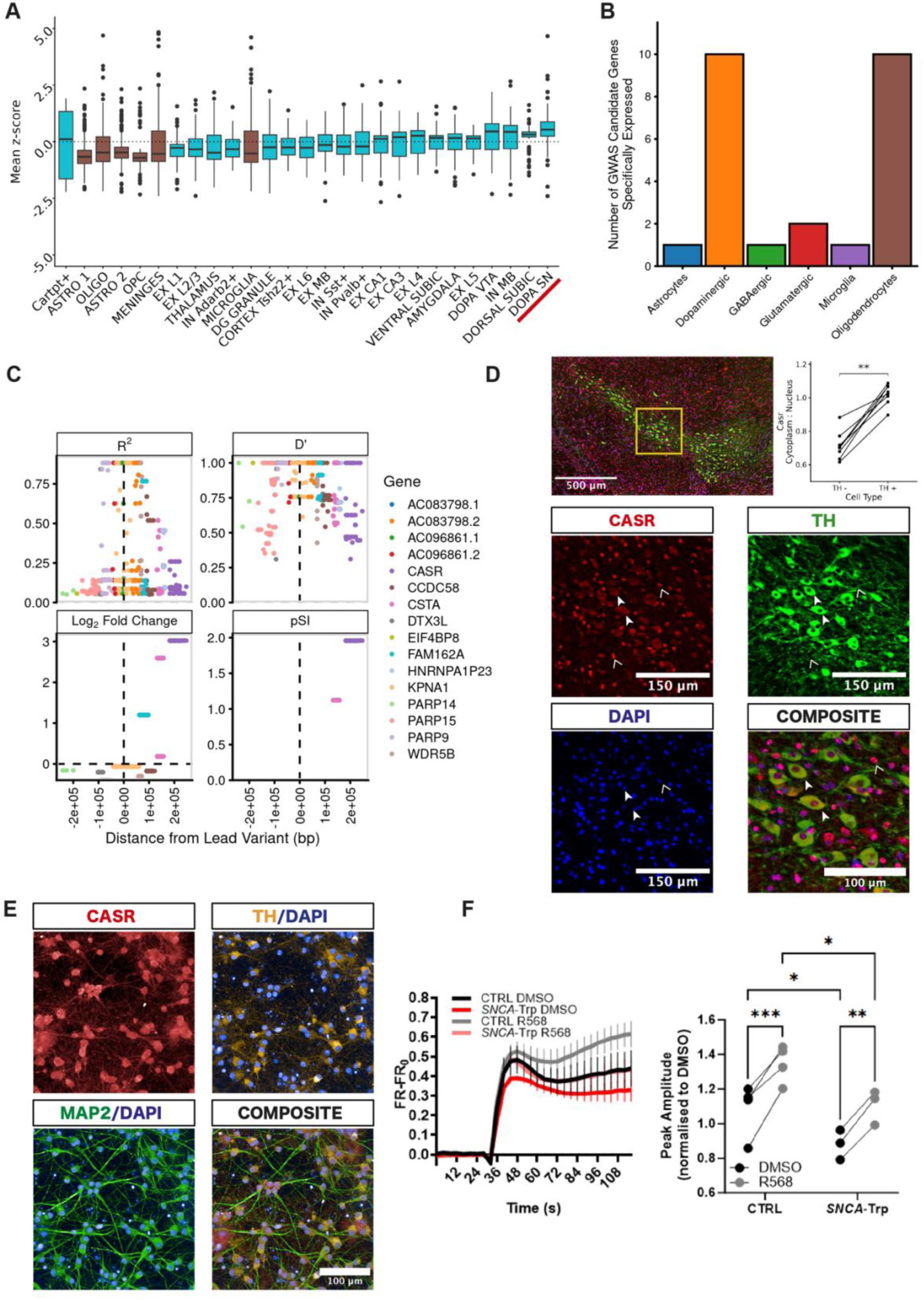
Mapping, expression, and function of PD GWAS risk loci. A) Log_2_-fold enrichment or depletion of candidate PD GWAS genes in each annotated cell type from Stereo-seq data. Genes were, with exceptions, generally enriched in neurons and depleted in glia. B) The number of candidate PD GWAS genes found to be significantly specifically expressed in each TRAP/RiboTag cell type tested. Dopaminergic neurons and oligodendrocytes jointly specifically expressed the greatest number of candidate genes (*n* = 56 (DAT-TRAP samples), 6 (GABAergic neurons), 6 (Glutamatergic neurons), 3 (Oligodendrocytes), 6 (Microglia), 6 (Astrocytes)). C) Linkage disequilibrium measures (R^2^ and D’) for rs55961674, indicating the candidacy of > 8 genes for consideration. There is significant linkage disequilibrium between the lead SNP and neighboring loci. Log_2_-fold change and pSI values indicate enrichment specificity to DA neurons. CASR was the most enriched and specifically expressed candidate gene. D) Immunohistochemical confirmation of specific Casr expression in TH-positive cytoplasm of the mouse ventral midbrain (filled arrows). Casr could be detected in neighboring nuclei of TH-negative cells (open arrows). The cytoplasm:nucleus ratio of Casr signal was significantly greater in TH-positive cells; paired wilcoxon rank sum text, (*n* = 8, *P* = 0.0078). E) Immunocytochemical confirmation of CASR expression in TH-positive induced pluripotent stem cell-derived dopaminergic neurons. F) Confirmation of CASR-mediated regulation of intracellular Ca^2+^ handling: R568 (a positive allosteric modulator of CASR) increased the Fura2 peak amplitude response to ionomycin administration (*n* = 3 per genotype; 2 genotypes; two-way repeated measures analysis of variance (ANOVA) by treatment group and genotype; *F* = 92.8*, P* = 0.0006, each colored line in the left panel is a representative trace for each genotype:treatment group).

We sought to demonstrate how cell type-specific gene expression could be used to prioritize candidate genes containing variation in linkage disequilibrium with lead PD SNPs. We observed that for 63 out of 91 testable PD GWAS loci, the lead SNP localized to a gene not considered specific or enriched in DAT-TRAP samples. Notable exceptions included rs356182, rs356203, rs256228, rs5019538 (*SNCA*), rs620513 (*FGF20*), rs11158026 (*GCH1*) and rs649339 (*SYT17*), in which the most proximal gene was also the most significantly enriched and specifically expressed. We focused on risk variant rs55961674, an intronic variant of *KPNA1*. In DAT-TRAP samples, *Kpna1* was depleted and in low abundance, relative to bulk ventral midbrain homogenate RNA, indicating low, nonspecific expression in DA neurons. Three candidate genes within the rs55961674 search window were significantly enriched by TRAP, specifically expressed (compared to other TRAP/RiboTag datasets) and contained variants in linkage disequilibrium with the lead SNP (Figure 3C). *Casr*, encoding the calcium sensing receptor, was selected for further investigation, based on demonstrating the most specific expression to DA neurons.

We first validated specific Casr protein expression in DA neurons of the mouse ventral midbrain (Figure 3D): Intriguingly, we observed a distinctly cytoplasmic pattern of expression specifically in DA neurons, while neighboring cells showed depleted cytoplasmic signal and intense nuclear staining. To confirm this difference, we compared the ratio of cytoplasmic to nuclear Casr intensity and found significantly higher cytoplasmic expression in TH-positive than TH-negative cells of the ventral midbrain (Figure 3D, Methods).

We next sought to investigate CASR expression and function in human pluripotent stem cell (iPSC)-derived DA neurons. We observed that CASR protein was prominently expressed in TH-positive neurons, with a diffuse cytoplasmic signal (Figure 3E). To evaluate whether CASR protein was functional and able to modulate intracellular calcium levels and dynamics in iPSC-derived DA neurons generated from a Parkinson’s patient carrying an *SNCA* triplication mutation, we measured cytoplasmic calcium in cells treated for one hour with R568 (10 μM), a positive allosteric modulator of CASR^44^. To estimate the levels of calcium stored in the intracellular compartments, we stimulated the cells with ionomycin (5 μM) and measured the increase in cytoplasmatic calcium using Fura-2AM^45^. A significant difference in ionomycin-evoked calcium release was observed between Parkinson’s and control iPSC-derived DA neurons before treatment with R568 (Figure 3F). Acute treatment led to increased evoked calcium release in both genotypes, bringing the patient-derived neurons into the range of the healthy control baseline samples (Figure 3F).

By integrating expression specificity data, GWAS summary statistics and the ability to functionally study gene function in iPSC-derived DA models, we can identify genes with novel function with a putative disease role in DA neurons.

### Stereo-seq captures age- and disease-induced expression changes across distinct cell types, loss of nigral dopaminergic neurons and neuroinflammatory expansion of microglia

Age remains the most important risk factor for neurodegeneration. We therefore examined aging-related changes in expression across all 29 cell types identified by Stereo-seq (Figure 4A). A range of cell types demonstrated differential expression (FDR-adjusted *P* < 0.05); most notably oligodendrocytes, cortical excitatory neurons of layers 1 to 3, thalamic neurons and astrocytes. Pathway enrichment analysis showed that a range of biological processes were affected, including axon ensheathment (oligodendrocytes), regulation of synaptic transmission and intracellular calcium ion homeostasis (excitatory cortical neurons) and regulation of catecholamine secretion (astrocytes).

**Figure 4:**
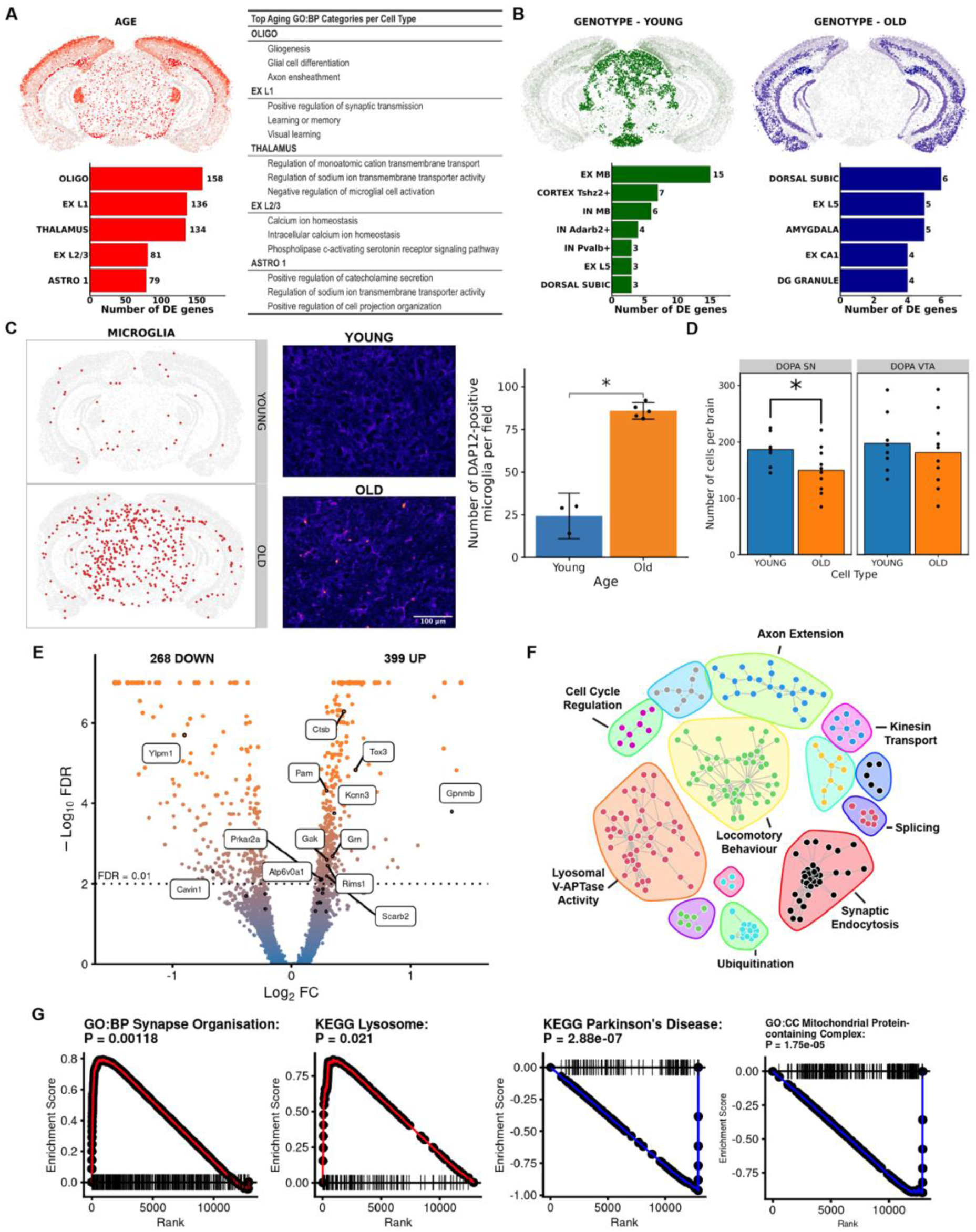
High resolution age-related changes in gene expression. A) Spatial representation of cell types ranked by overall aging-associated differential expression. Pathway overrepresentation analysis of each cell type revealed multiple aging-relevant processes. B) Spatial representation of cell types ranked by overall genotype-associated differential expression, stratified by age group. The magnitude of differential expression is subtler, compared to age. C) Odds ratio analysis of cell type abundance with age in Stereo-seq annotated cell types revealed the expansion of microglia in midbrain, corpus callosum and the external capsule. Confirmatory immunohistochemical staining for Tyrobp/Dap12, the leading transcriptomic marker of this cell population confirmed an increase microglial count in aged brain sections. Wilcoxon rank sum test, (*n* = 3 (Young brains), 5 (Old brains), *P* = 0.036, data here are represented as the mean ± 99 % confidence interval). Representative fields from young and old midbrain sections stained for Dap12/Tyrobp (Methods). D) Odds ratio analysis of DA neuron abundance in SN and VTA confirmed SN-specific loss of neurons with age. E) Volcano plot of the fold change and FDR of genes comparing aged and young TRAP samples (*n* = 56). 13 candidate PD GWAS (labeled) genes were within the genes significantly differentially expressed. F) Protein-protein interactions among TRAP-differentially expressed genes were used to identify functionally distinct clusters. G) Gene set enrichment analysis identifies and confirms the up- or downregulation of pathways relevant to aging in dopaminergic neurons. Genes related to synaptic organization and lysosomal biology were upregulated with age, as indicated by the enrichment score.

We next compared gene expression between *SNCA*-OVX and wild-type brains, stratified by age group (Figure 4B). In TRAP samples, overexpression of *SNCA* could be confirmed, on a background of normal *Snca* abundance (Supplementary Figure 4C). No other genes were identified as significantly differentially expressed in DA neurons between control and *SNCA*- OVX by TRAP. In Stereo-seq, changes were detected across a range of cell types (FDR-adjusted *P* < 0.05); in particular, excitatory and inhibitory midbrain neurons, cortical excitatory neurons of layer 5 and CA1 neurons of the hippocampus. Among the most significantly differentially expressed genes were *Ywhah*, *Ahcyl1*, *Nsamt1* and *Usp2*, each implicated in PD-relevant areas of biology (Ahycyl1 interacts with Tau, Nsamt1 plays a role in essential tremor, and *Usp2* deficient mice display altered locomotor activity).

By capturing individual cells, Stereo-seq enabled the comparison of cell number per cell type between conditions. We detected an expanded microglial population in aged brains (Figure 4C, FDR-adjusted *P* < 0.1). The expansion of activated microglia in aged brains was restricted to the midbrain, corpus callosum and external capsule. Immunohistochemical staining for Dap12/Tyrobp, a marker of microglia (and the most enriched Stereo-seq marker of this population) confirmed an increase in the number of Dap12-positive cells with a microglial morphology. We also observed a SN-specific loss of DA neurons with age, highlighting the vulnerability of this subpopulation of neurons compared to VTA (Figure 4D).

### TRAP reveals the extent of age-induced expression changes in dopaminergic neurons

We used TRAP to provide a deeper focus on DA neuron differential gene expression with aging. Greater measurement sensitivity and cohort size led to the discovery of 667 genes with altered expression in aged TRAP samples (399 upregulated, 268 downregulated, FDR-adjusted *P* < 0.01) (Figure 4E). Thirteen candidate PD GWAS genes were among the differentially expressed genes, including *Gpnmb*, also found to be enriched in both DA neurons and oligodendrocytes, and in agreement with previous age-related neuronal findings^10^. We used 1,931 high-confidence protein-protein interactions to subdivide differentially expressed genes into functionally related clusters (Figure 4F, Methods). Each cluster of genes was distinctly enriched for terms related to neuronal function, with particular importance in PD (e.g. Lysosomal V-ATPase activity, locomotory behavior and synaptic endocytosis). By taking the directionality of expression change into account, gene set enrichment analysis further indicated an upregulation of synaptic and lysosomal-related genes (e.g. *Syt1, Syt11, Sv2a, Ap2b1*, *Atp6v0a1, Atp6v0d1, Atp6v1e1, and Atp6ap1*) and a downregulation of mitochondrial and PD-related genes (*mt-Co1, mt-Nd1, mt-Nd4, mt-Nd5, mt-Cyb*) (Figure 4G).

## Discussion

In this study, we generated a single cell-level spatial transcriptomic map of gene expression in the adult mouse brain and produced a high-fidelity translatome-level profile of DA neuron gene expression. By integrating these two data modalities, we characterized the distinctive expression features of DA neurons, demonstrated how expression specificity can be used to prioritize candidate causal genes in PD, and examined the changes that occur in the brain and DA neurons specifically with age.

We identified 29 distinct cell types by unsupervised clustering of cells, based on their expression properties. The spatial compartmentalization of distinct cell types could be mapped to anatomical regions, as in the case of neuronal populations of the hippocampus, thalamus, and midbrain. The ability to spatially visualize each cluster aided the annotation of cell types, however as methods of spatial transcriptomic analysis develop, we anticipate the integration of spatial information directly into the clustering process, wherein cells with close spatial proximity or patterned localization (e.g. cortical, intestinal and dermal layers) and matched expression are more considered more similar.

The capture area of each Stereo-seq array is 100 mm^2^ in this study; larger than existing available spatial transcriptomic technologies (Visium, 42.25 mm^2^; Slide-Seq 7.1 mm^2^)^46–48^, and each Stereo-seq array contains 40 billion capture spots (Visium, 5,000; Slide-Seq, tens of thousands). We leveraged this greater size and density of information to spatially resolve individual cells across the brain. By achieving nanoscale resolution, single-cell spatial data were generated without the need for deconvolution-based methods. We also did not require matched single-cell RNA-seq samples for cell identity annotation.

By integrating Stereo-seq and TRAP data, we identified the expression features most specific to DA neurons in comparison to neighboring cell types. While Stereo-seq provided spatial context, single-cell resolution, and a greater number of unique cell types for comparison, TRAP demonstrated greater measurement sensitivity. By integrating short- and long-read sequencing data from TRAP RNA, we have been able to profile the state of splicing in DA neurons, revealing differential transcript usage across more than a thousand genes. We detected 817 instances of alternative splicing in which individual isoforms were enriched by TRAP, but the overall gene-level count was not. This finding indicates that mouse DA neurons actively translate a larger number of genes than are detectable from only gene-level count data. Our understanding of cell type-specific expression is strengthened by considering transcript-level expression data.

The single-cell resolution of Stereo-seq enabled the distinction of SN and VTA DA neurons in ventral midbrain. Although we detected subtypes within both populations, we have not reported them here, as their identity could not be accurately predicted by gene expression in testing data. We expect the addition of further Stereo-seq samples, coupled with improved spatial clustering method development will lead to the robust identification of further DA neuron subtypes in our data.

We used measures of expression specificity from Stereo-seq and TRAP to demonstrate how genes within disease risk loci can be prioritized for investigation. We showed that candidate causal gene expression was most specific to SN DA neurons, supporting the hypothesis that selective vulnerability may be in part conferred by the selective expression of causative genes^41^. We focused on a window surrounding rs55961674 for study, as this SNP falls within the intronic region of *KPNA1*; a gene considered to not be expressed abundantly or with specificity in DA neurons. By leveraging the sensitivity of TRAP and the diversity of cell types detected in Stereo-seq, we could rank linkage-associated genes surrounding rs55961674 by their expression specificity. We demonstrated specific dopaminergic cytoplasmic expression of CASR in two model systems (mouse and human), and showed that CASR regulates intracellular calcium handling in human DA neurons. The specificity of cytoplasmic CASR expression to DA neurons and demonstration of its functional role in regulating intracellular calcium handling indicates that variation in this gene could contribute to PD disease risk. Long-term functional study of this gene is of interest, such as by generating a DA neuron-selective knockout *Casr* mouse model.

The large capture area of the Stereo-seq array, combined with single cell-level resolution, enabled parallel region-dependent and cell type-dependent comparison. We identified an age-dependent reduction in SN DA neuron cell number. This supports previous findings that DA neuron number declines with age^1,2^ and demonstrates the ability of Stereo-seq to identify a subtype of DA neuron that is most vulnerable to aging. In addition, we detected and confirmed the expansion of microglia with age, expressing a variety of pro-inflammatory markers. Previous investigations into changes in microglial number with age have reported disparate findings, depending on the species and brain region investigated^10,49,50^. However, age-related neuroinflammation has been shown previously to significantly contribute to degeneration in PD^51^.

A surprising finding in this study was the absence of detectable gene expression changes in DA neurons due to *SNCA* overexpression. The age-dependent pattern of SN DA neuronal loss in *SNCA*-OVX mice is considered to recapitulate the slow progression of PD pathology in patients. We would suggest that the transcriptional effects of *SNCA* overexpression (at the level achieved by the *SNCA-*OVX model) are subtle. To capture DA neuron-specific gene expression changes induced by the interaction of *SNCA* overexpression and aging, it may be necessary to sample a greater number of timepoints and replicates. In addition, the *SNCA*-OVX mice used in this study also expressed *Snca* (Methods). The expression of wild-type *Snca* may ameliorate disease-related pathology previously observed in *Snca^-/-^ SNCA*-OVX mice. However, our Stereo-seq analysis did enable the detection of transcriptional changes due to *SNCA* overexpression in other cell types.

We leveraged the sensitivity of TRAP to assess the evidence for axonal translation in DA neurons. We observed enrichment of DA markers, *Th, Slc6a3 (DAT), and Slc18a2 (VMAT2)* in striatal TRAP samples, however we also observed enrichment of markers of other cell types. Furthermore, the abundance of DA markers in striatal TRAP samples was markedly lower than in midbrain-derived samples. We detected GFP puncta in striatum that colocalized with TH, although overall signal was sparse.

Hobson et al. previously used RiboTag, a similar technology to TRAP, to study axonal translation in DA neurons and concluded that there was no evidence for the process occurring. We conclude that we do see evidence for axonal translation in DA neurons by TRAP, although we suggest that the scale of activity is substantially lower than at the level of the cell body. The enrichment of markers of other cell types in striatal TRAP samples also indicates the likelihood of low-level ectopic expression of the eGFP-L10a transgene in neighboring cells, as has been reported in DAT-Cre lines^52^.

### Limitations of the study

An ongoing challenge for droplet-based single cell and emerging spatial transcriptomic technologies will be to increase the sensitivity of gene detection. Lower detection sensitivity leads to zero inflation, in which many genes are detected at the population level but are largely undetectable at the individual cell level^53,54^. The power to detected differential expression is limited by zero inflation. Lateral diffusion of RNA also limits the resolving capacity of current spatial transcriptomic methods^55^. To minimize the impact of diffusion, we segmented individual cells by spatial read intensity, excluding lower intensity background regions. In addition, we excluded all cells that co-expressed markers of neurons and glia. TRAP demonstrated greater detection power (over 15,000 genes per sample), offering the most inclusive and sensitive measure of DA gene expression currently available. The limitation of TRAP is its specificity to cell subpopulations. DA neurons of the SN and VTA could not be reliably separated during tissue processing due to their proximity, so measurements reflect the averaged expression of these regions. With greater detection sensitivity and/or sample number, further DA neuron subpopulations may become identifiable.

Together, our spatial transcriptomic and translatome profiling of DA neurons represent a valuable resource to the neuroscience community. Our spatial data can be used to prioritize candidate causal genes involved in conferring genetic risk of brain-related diseases other than PD. In addition, these data can be used as a reference for the development of novel analytical approaches to spatial research. Our TRAP data provides a reference for the querying of genes or isoforms specific to DA neurons in health and with age. We have combined all results from analyses in this study into a database for public access: spatialbrain.org

## Methods

### Ethics statement and animal husbandry of Mus musculus

All experiments and procedures conducted on animals were carried out in accordance with United Kingdom Home Office regulations under the Animal (Scientific Procedures) Act (1986), and were approved by the local ethical review board of the Department of Physiology, Anatomy and Genetics at the University of Oxford. All mice were housed in the University of Oxford Biomedical Services Building. Mice had access to standard food and water ad libitum. Mouse holding rooms were maintained at 22 °C and 60 to 70% humidity on a 12-hour light-dark cycle. All mice were bred on a C57/BL6 background (MGI Cat# 2159769, RRID:MGI:2159769). For all experiments, animals of both sexes were used unless indicated otherwise. Animals used for both Stereo-seq and TRAP technologies were litter mates.

### Transgenic model generation

Homozygous Rosa26^fsTRAP^ mice were bred with homozygous DAT^IREScre^ mice to generate heterozygous Rosa26^fsTRAP^::DAT^IREScre^ offspring that were used for TRAP experiments. Rosa26^fsTRAP^ parents were crossed and bred to be hemizygous *hSNCA+* (*SNCA*-OVX): Offspring were a mixture of hemizygous *hSNCA+* and *hSNCA-.* All animals were wild-type mouse *Snca^+/+^*, due to the position of the ROSA26-eGFP-L10a locus being on the same chromosome as *Snca* (chromosome 6). Routine ear-clipping, digestion and PCR-genotyping was performed as described in Hunn et al^7^. Primer sequences used for confirmation of transgene expression are supplied below^8,9,34,35^.

### TRAP

Mice were sacrificed by cervical dislocation, the ventral midbrain and striatal tissue were rapidly dissected and affinity purification of eGFP-tagged polysomes was performed as previously described^20^. Briefly, dissected tissue was immediately dounce homogenized in lysis buffer containing 20 mM HEPES KOH, 150 mM KCl, 10 mM MgCl2, 0.5 mM DTT, 100 µg/mL cycloheximide, RNasin (Promega) and SUPERase-in (Life Technologies) and Complete-EDTA-free protease inhibitors (Roche) in Rnase-free water. The lysate was cleared by two-stage centrifugation for 10 minutes at 2,000 x *g* and 20,000 x *g*. Each lysate was incubated with monoclonal anti-GFP antibodies (Heintz Lab; Rockefeller University Cat# Htz-GFP-19C8 and Htz-GFP-19F7, RRID:AB_2716737 and RRID:AB_2716736), coated on paramagnetic beads through a streptavidin-biotin-protein L linker (Pierce; Thermo Fisher Scientific) for 18 hours at 4°C. To remove non-specifically bound material, including RNA, beads were washed 6 times in a high-salt solution containing 20 mM HEPES KOH (pH 7.4), 350 mM KCl, 10 mM MgCl2, 1% NP-40, 0.5 mM DTT, 100 g/ml cycloheximide, and Rnasin and Superasin. RNA was extracted using the Rneasy Micro Plus kit (Qiagen). RNA quantity and integrity were measured using the Quant-it™ RiboGreen RNA Assay Kit (Thermo Fisher) and Agilent 2100 Bioanalyzer, respectively. RNA-seq library preparation was performed using the NEBNext Ultra II Directional RNA kit. Transcript level quantification was performed using Salmon (v1.4.0, RRID:SCR_017036)^56^.

### Long-read sequencing and data processing

Twelve TRAP samples and three TOTAL samples were sequenced using the Oxford Nanopore Technologies MinION platform. TRAP samples were equally divided by age and genotype (N = 3 per age:genotype). Library preparation was performed using the cDNA-PCR kit (SQK-PCS109). Raw fast5 data was basecalled and demultiplexed using Guppy (v4.5.2). Read data from FASTQ files were aligned to the mm10 genome (Gencode M25 GRCm38.p6) using minimap2 (v2.18, RRID:SCR_018550)^57^. Transcript level quantification was then performed using Salmon (v1.4.0, RRID:SCR_017036)^56^.

### Stereo-seq library preparation and sequencing

Stereo-seq libraries were prepared as previously described by Chen *et al*. (2022), using one chip per brain, across 18 brains in total. In brief, Stereo-seq samples were first prepared by collecting postmortem mouse brains and flash-freezing at -80°C. 10 μm tissue sections were collected -3.5 mm from bregma (to optimize capture of DA neurons in the SN and VTA) using a Leica CM1950 cryostat and adhered to each Stereo-seq chip (BGI Research). The chip was placed on a warming plate at 37°C for 3 minutes and fixed in methanol at -20°C for 30 min. The chip was incubated with 100 μL 0.1 % pepsin at 37°C for 12 min for permeabilization and washed with 0.1× SSC buffer containing 0.05 U/μL RNase inhibitor. RNA captured by the DNA nanoball on the chip was reverse transcribed at 42°C for 90 min. Tissue was removed from the chip by incubating with tissue removal buffer at 55°C for 10 min. After washing with 0.1× SSC buffer, the chip with cDNA was incubated with 400 μL cDNA release buffer at 55°C for 4 hours. cDNA was purified and amplified using cDNA primer. A total of 20 ng of cDNA was fragmented, amplified, and purified to generate each cDNA sequencing library. The cDNA library was sequenced on a MGI DNBSEQ-Tx sequencer with the read length of 50 bp for read 1 and 100 bp for read 2.

### Immunohistochemistry

Mice were anaesthetized by intraperitoneal injection of 150 µl pentobarbital. Upon loss of the pedal withdrawal reflex, the thoracic cavity was opened, and 25 mL of phosphate-buffered saline (PBS) was administered transcardially, followed by 25 mL of 4% paraformaldehyde (PFA), diluted in PBS. The brain was extracted and stored overnight in 4% PFA at 4 °C. Brain tissue was next dehydrated, and paraffin embedded through graded alcohols, histoclear (ThermoFisher Scientific) and paraffin solution. Tissue blocks were finally sectioned at 7 µm using a Leica microtome (Leica Microsystems, Wetzlar, Germany) and mounted on glass slides (VWR, Superfrost® Plus).

Sections were prepared for immunofluorescence by performing dewaxing in histoclear, rehydration in graded alcohols, antigen retrieval by microwave heat (20 min) and citrate buffer (ab96678, Abcam). Tissue was blocked for 1 hour at room temperature using 10% donkey serum in TBS. Primary antibodies to TH (Abcam Cat# ab76442, RRID:AB_1524535, 1:500), GFP (Thermo Fisher Scientific Cat# A-11122, RRID:AB_221569, Invitrogen, 1:1000), CASR (Abcam Cat# ab79038, RRID:AB_2071489, 1:100) and TYROBP/DAP12 (Abcam Cat# ab283679, 1:100) were prepared in blocking solution and incubated overnight at 4 °C. Slides were washed using TBS and incubated in secondary antibodies (Alexa Fluor-conjugated anti-rabbit/sheep 1:1000) and DAPI (Thermo Fisher Scientific Cat# D3571, RRID:AB_2307445, 1:5000) for 1 hour at room temperature. Slides were washed and coverslips mounted using FluorSave mounting medium (Millipore, Massachusetts, United States of America).

For detection and quantification of Tyrobp/Dap12-positive cells and analysis of Casr-positivity, two custom pipelines were built using CellProfiler, provided in the accompanying GitHub repository to this manuscript^58^. To calculate the Casr cytoplasm:nucleus ratio, a 10 pixel-wide cytoplasmic object was created, originating from each DAPI-positive spot (nucleus). Each nucleus and cytoplasm was classified either as TH-positive or TH-negative, based on overlapping cytoplasmic TH signal. Objects with only partial TH overlap were filtered, to ensure accurate classification. The maximal Casr intensity for each object was recorded and summarized for each mouse by calculating the mean across objects. The Casr cytoplasm:nucleus ratio represents the ratio of these means.

### Immunocytochemistry

After R568 treatment, the iPSC derived DA neurons were fixed in 4% PFA. Cells were permeabilized for 10 min a solution of 5% NDS, 1% BSA and 0.5% Triton-X 100. Samples were blocked in a solution of 5% NDS and 1% BSA for 1 h. Cells were then incubated for 16 h O/N at 4°C with the following primary antibodies: CASR (Abcam Cat# ab79038, RRID:AB_2071489, 1:500), MAP2 (Abcam Cat# ab92434, RRID:AB_2138147, 1:1000), tyrosine hydroxylase (Millipore Cat# AB1542, RRID:AB_90755, 1:500). Cells were washed twice in PBS and then incubated in secondary antibodies and DAPI for 1.5 h at RT. Cells were washed 3 times and stored in fresh PBS until imaging. Cells were imaged using the OperaPhenix (PerkinElmer).

### Differentiation and culturing of iPSC-derived dopamine neurons

Three control and three SNCA-Triplication patient lines underwent a differentiation process following the protocol described by Fedele et al.^59^ with slight modifications as outlined in Williamson et al.^60^. Briefly, the cells were initially patterned for 10 days, followed by expansion of midbrain floor plate progenitors for 19 days. Subsequently, the progenitors were differentiated for an additional 10 days, then replated and matured for a further 5 weeks until reaching DIV 60 for imaging.

### Fura-2

Fura-2 AM was prepared to 5 mM concentration in calcium-free HBSS supplemented with 20 mM HEPES. It was then diluted to the final concentration of 2.5 mM in Neurobasal medium supplemented with B27 and L-glutamine, with DMSO (0.1%) or the CaSR positive modulator R568 (10 μM).

iPSC-derived DA neurons were incubated in the solution of Fura-2 AM and drug for 1 h at 37°C, 5% CO_2_ and then imaged on a FlexStation 3 Multi-Mode Microplate Reader (Molecular Devices) at 37°C. The dye was excited at 340 nm and 380 nm and was detected at 510 nm. Each well was imaged every 4 s for 100 s and injected with ionomycin (final concentration 5 μM) after a baseline of 28 s.

For the analysis, the 340/380 ratio was computed, and then the baseline was subtracted from all the timepoints, to obtain a normalized trace for each well. The maximum intensity (peak amplitude) of the normalized trace was found and the area under the curve (AUC) was calculated using the left rectangular approximation method.

### Processing of raw Stereo-seq data, quality control and cell type identification

Raw gene-by-spot data per sample were aggregated to create a 2-dimensional image of RNA signal for each sample using custom Python (v3.9.0, RRID:SCR_008394) scripts (Supplementary Materials). To segment individual cells, each image was subjected to a processing pipeline written in Python (workflow illustrated in Supplementary Figure 1). Steps taken: **Mask generation from unspliced counts:** Gaussian filter (sigma = 5), background subtraction (white tophat, 50 pixels), Otsu thresholding, conversion to mask, fill holes, watershed. **Mask generation from spliced + unspliced counts:** As for unspliced counts. Watershed boundaries from the spliced + unspliced mask were then subtracted from the unspliced mask. Objects were retained from the spliced + unspliced mask that overlapped with the unspliced mask. Each cell was labelled using the label function in SciPy (v1.9.0, RRID:SCR_008058). Raw gene-by-spot data were then aggregated to the gene-by-cell level and imported into scanpy (v1.9.1, RRID:SCR_018139)^61^.

The initial Stereo-seq dataset contained 497,766 cells. In a first round of filtering, low complexity and putative doublet/triplet cells were filtered. To remove low complexity cells, a minimum gene detection cutoff of 200 was selected. To remove putative doublet/triplet cells, a maximum gene detection cutoff was set on a per-brain basis to the median number of genes detected + 4 median absolute deviations. After first round filtering, 415,402 cells remained. Count data were subsequently processed using SCVI, including sample brain of origin as a categorical covariate, and the number of genes detected per cell as a continuous covariate^62^. A uniform manifold projection (UMAP) was generated for cells from each mouse brain and overlaid, showing a similar pattern of separation (Supplementary Figure 1A).

Cell type identification was performed in two rounds. In a first pass, leiden clustering was performed at iteratively greater resolutions^63^. Cell types without clear spatial organization e.g. some glial types, were annotated based on marker gene enrichment. Remaining cells were then processed using SEDR in order to include spatial information in the clustering process^64^. Mclust was used for clustering spatially defined cell types^65^. If a cluster contained fewer than 200 cells, it was considered final and no further subclustering was performed. Dopaminergic neurons were labelled according to their anatomical region (SN and VTA), based on their spatial coordinates.

For differential expression analysis in Stereo-seq samples, gene counts were normalized to the total number of counts per cell and log-transformed. Marker genes for each cluster were identified using the Wilcoxon rank-sum test, providing two-sided *P* values. The identity of each cell type was annotated by integrating marker gene data with previous literature and by confirming the spatial distribution of clustered cells.

A second round of cell filtering was performed after clustering to remove cells that were enriched in both *Snap25* and *Plp1*. We suspected that these cells represent neuronal/glial contaminated mixtures, as previously reported in single cell data^66^. After second-round filtering, 355,307 cells remained.

### Spatially variable gene detection

Spatially variable genes were identified using SpatialDE2 with default settings. Coordinates and full expression matrices of each brain were supplied, divided by cell type. To combine results from separate samples, a meta-analysis was performed on the raw *P* values from each sample, using Fisher’s method. The Benjamini & Hochberg correction for multiple comparisons was subsequently used and genes were considered spatially variable with an adjusted *P* value of < 0.01.

### Cell type abundance analysis in Stereo-seq data

To test for differential abundance of cell types between age groups, we used mixed effects modelling of associations of single cells (MASC). MASC tests whether cell type membership of individual cells is influenced by an experimental covariate of interest, while accounting for technical covariates and biological variation. We specified mouse genotype as a fixed covariate and mouse of origin as a random effect in the generalized, mixed-effect model. Changes in cell type abundance were considered significant at FDR-*P* < 0.01.

### Differential gene expression analysis

Differential gene expression analysis in TRAP samples (including calculation of gene enrichment and depletion, relative to tissue homogenate RNA) was performed using DESeq2 (v1.36.0, RRID:SCR_015687) in R (v4.2.1, RRID:SCR_001905) with Bioconductor (v3.15, RRID:SCR_006442). Adaptive shrinkage of log fold change estimates was performed using ashr. The following settings were changed from defaults: minReplicatesForReplace = Inf, cooksCutoff = Inf, filterFun = ihw, lfcThreshold = log_2_(1.05). Genes were classed as significantly differentially expressed with an FDR-adjust *P* value < 0.01.Protein-protein interactions were obtained using STRINGdb (version 11) with a minimum confidence threshold of 0.4^67^.

### Specificity index calculation in TRAP/RiboTag samples

To calculate the specificity index of genes detected within TRAP/RiboTag datasets of dopaminergic neurons and other cell types within midbrain, pSI (v1.1) was used with default settings^22,24,38–40,68^. Genes were considered significantly specifically expressed with an FDR-adjusted *P* value < 0.01.

### Differential transcript usage analysis

Differential transcript usage was performed using DRIMSeq (v1.24.0). To consider only genes containing more than 1 isoform, with a minimum level of expression, the following parameters were used: min_samps_gene_expr = n * 0.75, min_gene_expr = 10, min_samps_feature_expr = 10, min_feature_expr = 10, min_samps_feature_prop = 10, min_feature_prop = 0.1.

### GWAS prioritization analysis

A list of 303 genes (sourced from Nalls et al., 2019 supplementary materials), containing SNPs at an r^2^ > 0.5 and located within ±1 Mb of 107 common risk variants for sporadic PD was used for prioritization analysis^42^. To convert between human and mouse gene symbols, homologene (v1.4.68.19.3.27, RRID:SCR_002924) and biomaRt (v2.52.0, RRID:SCR_019214) were used^69,70^. TRAP enrichment (measured as the product of the log_2_ fold-change and FDR-adjusted *P* value) and specificity indices for DAT-TRAP samples were used for gene prioritization, per lead SNP.

### Gene set enrichment analysis

Gene set enrichment analysis was performed using fgsea (v1.22.0, RRID:SCR_020938) with the following parameters: eps = 0, minSize = 10, maxSize = 500, nPermSimple = 10000. Pathway data were obtained from the Molecular Signatures Database (v7.5.1, RRID:SCR_016863), using C2 Canonical Pathways from Kegg and C5 Gene Ontology _sets71,72._

### Statistics and reproducibility

No statistical methods were used to predetermine sample sizes, but our sample sizes for TRAP analyses surpass those reported in previous publications^23,28,73^ and our sample sizes for Stereo-seq samples are comparable with those of similar spatial transcriptomic datasets^11^. All statistical analyses were performed with R (v4.2.1) and Python (v3.9). All *P* values were modified to an FDR of 1, 5 or 10 % as described in the text with the Benjamini & Hochberg method.

### Data and code availability

All raw data from TRAP experiment have been uploaded to the Gene Expression Omnibus (GEO accession number: GSE215276). The Stereo-seq raw data that supports the findings of this study have been deposited into CNGB Sequence Archive (CNSA) of China National GeneBank DataBase (CNGBdb) with accession number CNP0003397. All processed data, including results of all analyses are provided at spatialbrain.org. All code for processing data from raw input or processed counts are provided at https://github.com/peterkilfeather/spatialbrain. Protocols for laboratory collection and processing of samples as described in this study are published as a collection at protocols.io (DOI: Pending publication).

## Authors Contributions

P.K. performed all analyses, with help from J.H.K. and H.L. P.K. and K.W. optimized and performed TRAP experiments. P.K. performed all long-read sequencing experiments, all immunohistochemistry experiments, and all tissue collection for Stereo-seq. M.C.C. performed Fura-2 CASR and iPSC ICC experiments. Y.A., J.H.K., X.Z. and X.C. performed the Stereo-seq experiments. N.C.-R. co-supervised the project and contributed to project and experimental design. R.W.M conceived the project. R.W.M and Z.S. supervised the entire study and secured funding. P.K. and R.W.M. wrote the paper, with contributions from all authors. All authors reviewed the manuscript and approved its submission.

## Acknowledgements

This research was funded in part by Aligning Science Across Parkinson’s [ASAP-020370] through the Michael J. Fox Foundation for Parkinson’s Research (MJFF), in part by the Monument Trust Discovery Award from Parkinson’s UK (J-1403), and in part by Science, Technology and Innovation Commission of Shenzhen Municipality under grant No. JCYJ20180507183628543. We sincerely thank the support provided by China National GeneBank. For the purpose of open access, the author has applied a CC BY public copyright license to all Author Accepted Manuscripts arising from this submission. P.K. was supported by a Medical Research Council studentship and post-doctoral Fellowship; K.W. was supported by a Parkinson’s UK studentship (H-1301) and M.C.C. is supported by the Wellcome Trust (Grant ref: 223202/Z/21/Z) and previously held the Joan Pitts-Tucker/Heyman Moritz Studentship.

---

**Supplementary Figure 1:**
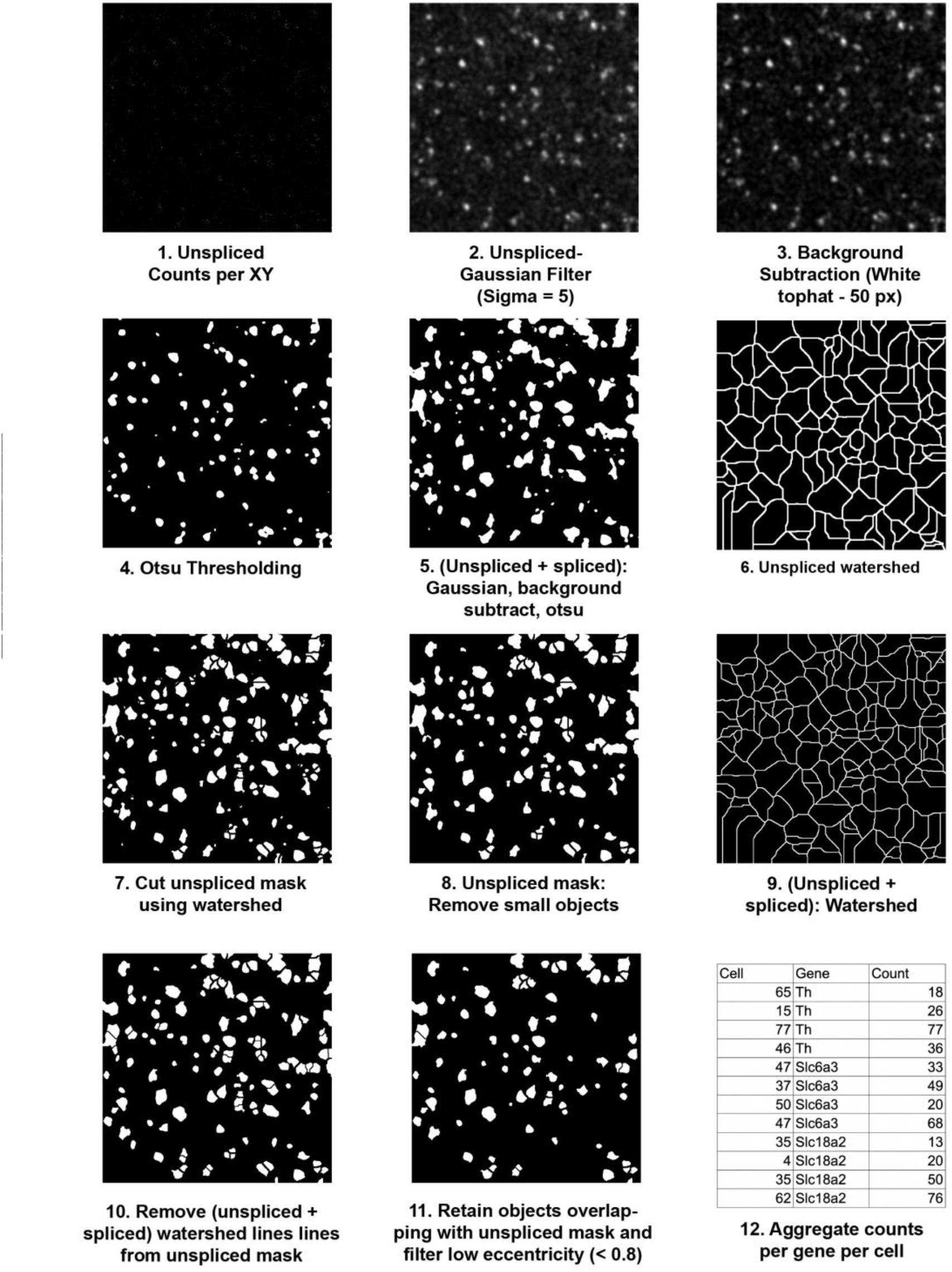
Stereo-seq segmentation pipeline. 1. A transcript “map” of unspliced reads was generated by calculating the total number of unspliced counts detected per spatial X-Y position across all genes. 2. Gaussian filtering (sigma = 5) to enhance signal from cell regions and reduce signal from the background. 3. White tophat background subtraction to further remove background signal. 4. Otsu thresholding to generate an unspliced mask. 5. Repeat of steps 1-5 using combined spliced and unspliced counts to generate a total count mask. 6. Watershed segmentation of the unspliced mask. 7. Subtraction of watershed boundaries from the unspliced mask. 8. Filtering to remove very small objects from the unspliced mask. 9. Watershed segmentation of the total count mask (unspliced + spliced). 10. Subtraction of the total count watershed boundaries from the unspliced mask. 11. Filtering to retain objects from the total count mask that overlap with objects from the unspliced mask. 12. Aggregation of gene counts to the cell-level.

**Supplementary Figure 2:**
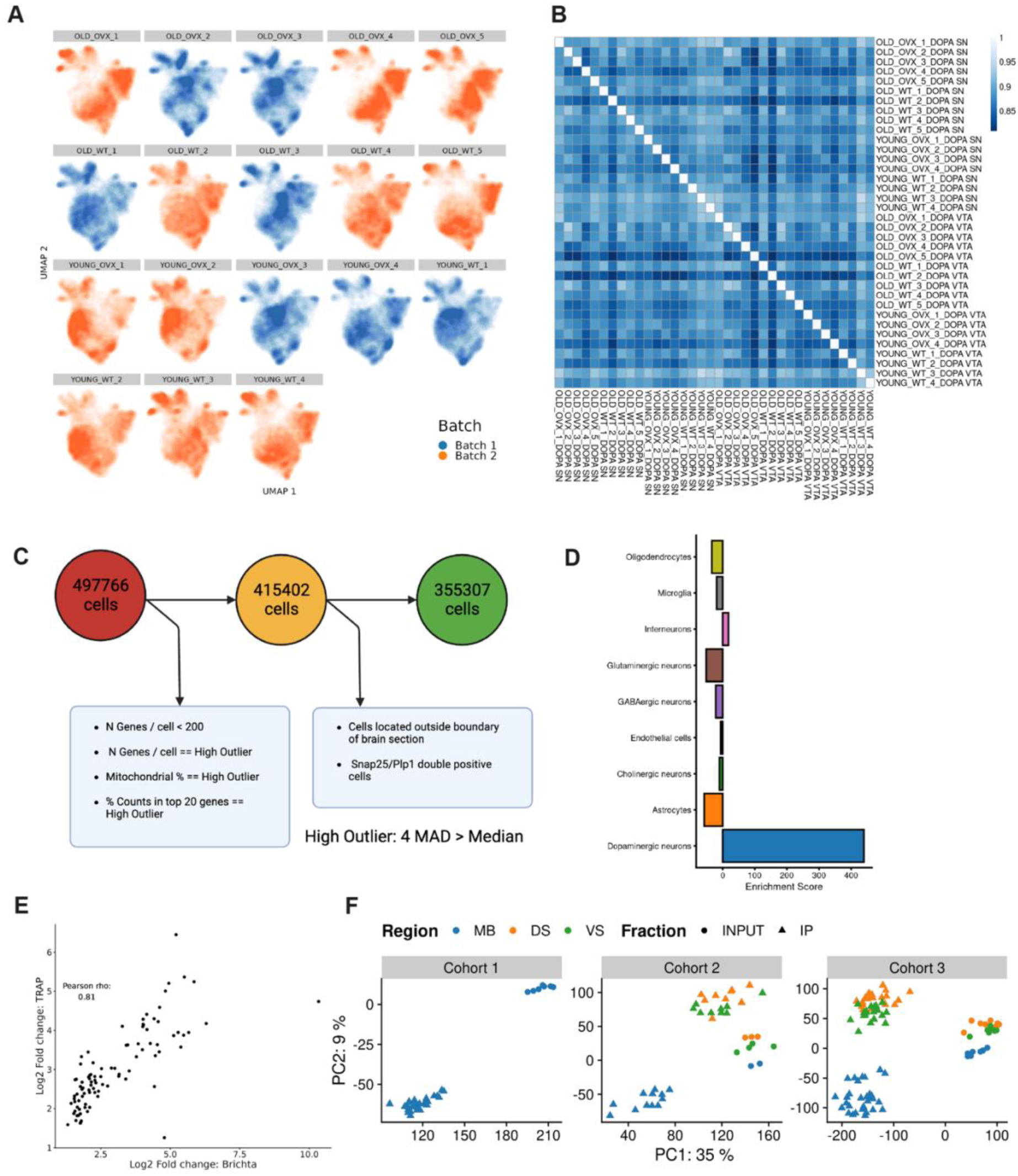
Stereo-seq and TRAP quality control. A) UMAP representation of all 18 brains, colored by age. No obvious batch, age or genotype-related differences in UMAP distribution were observed. B) Heatmap of sample-to-sample pearson correlation of pseudobulk dopaminergic neurons from SN and VTA. Raw counts were vst-transformed using DESeq2. C) The number of cells retained at each filtering step. During unsupervised clustering, a subset of cells contained both neuronal and glial markers (e.g. Snap25/Plp1). These cells were excluded from further analysis, as their identity could not be confirmed. D) Enrichment score (-log_10_ *P* multiplied by log_2_ fold-change) for cell types likely to be resident within dissected midbrain tissue for TRAP. DA neurons were the only cell type, out of 178 that showed clear enrichment. E) Principal component analysis (PCA) biplot of TRAP samples. TRAP samples were collected in three cohorts, as shown. Samples were most clearly distinguished along PC1 and PC2 by the effect of TRAP capture, separating TRAP and TOTAL homogenate samples and by the region of origin (MB, midbrain; DS, dorsal striatum; VS, ventral striatum).

**Supplementary Figure 3:**
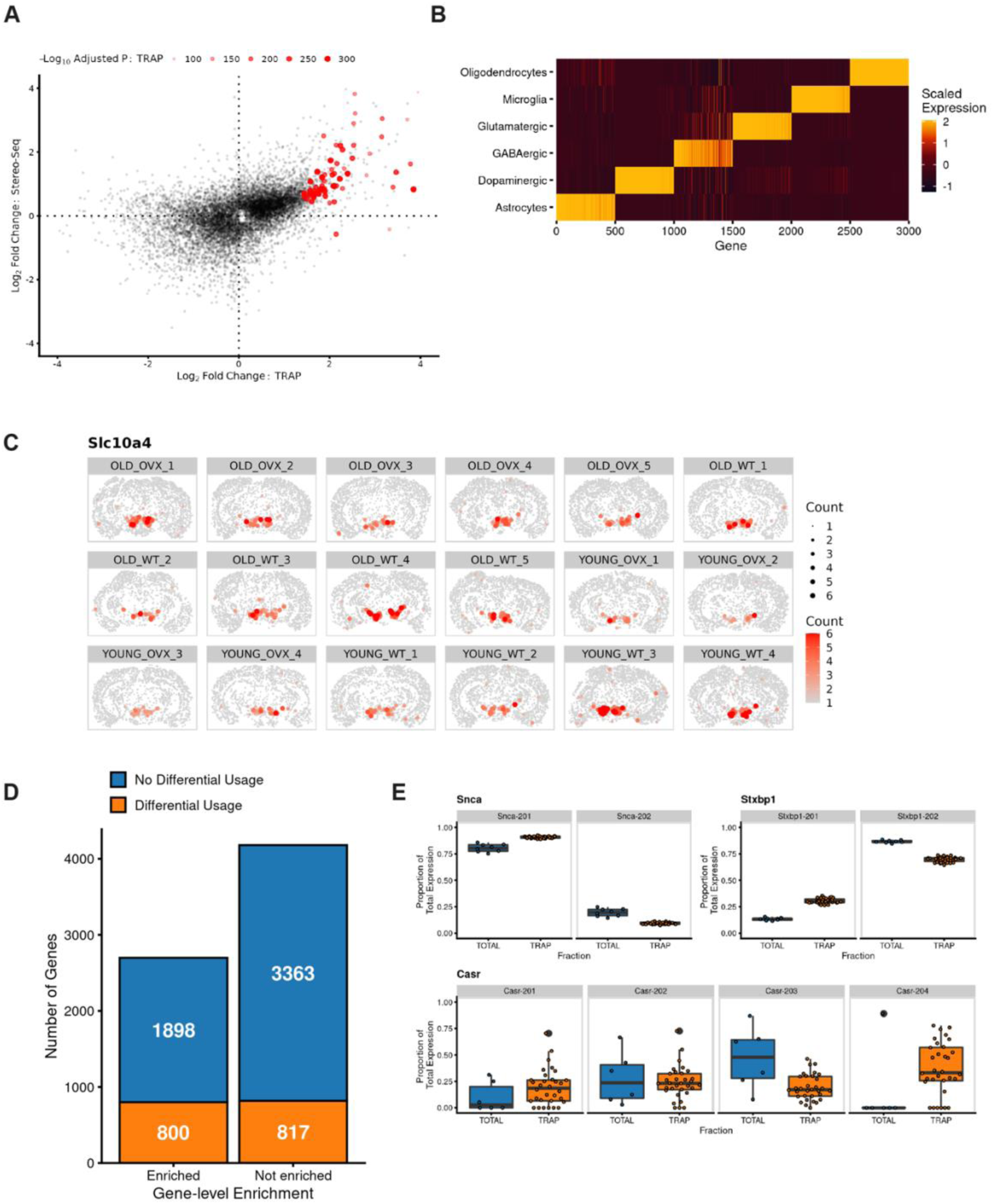
Specificity analysis of TRAP/RiboTag datasets and DA splicing summary. A) Log_2_ fold-change of genes detected in TRAP and Stereo-seq datasets between old and young samples. The concordance between both technologies is illustrated by the predominance of genes in the bottom left and top right quadrants. The top 100 TRAP-enriched genes are highlighted in red. B and C) Example of Slc10a4 demonstrating DA neuron enrichment in Stereo-seq brain sections. D) Scaled expression heatmap of the top 500 genes per cell type that are specifically expressed in each TRAP/RiboTag dataset. Yellow indicates high relative expression of a given gene in that cell type. Most genes are expressed in each target cell type more than 2 standard deviations above the expression level of all other cell types. E) Summary of the number of the genes found to exhibit differential transcript usage in midbrain TRAP samples, relative to midbrain homogenate RNA. Differential transcript usage is evident to an equal degree in genes enriched and not enriched by TRAP. F) Example of differential transcript usage of *Snca*<u>, *Stxbp1*, *Casr*</u> in midbrain TRAP samples.

**Supplementary Figure 4:**
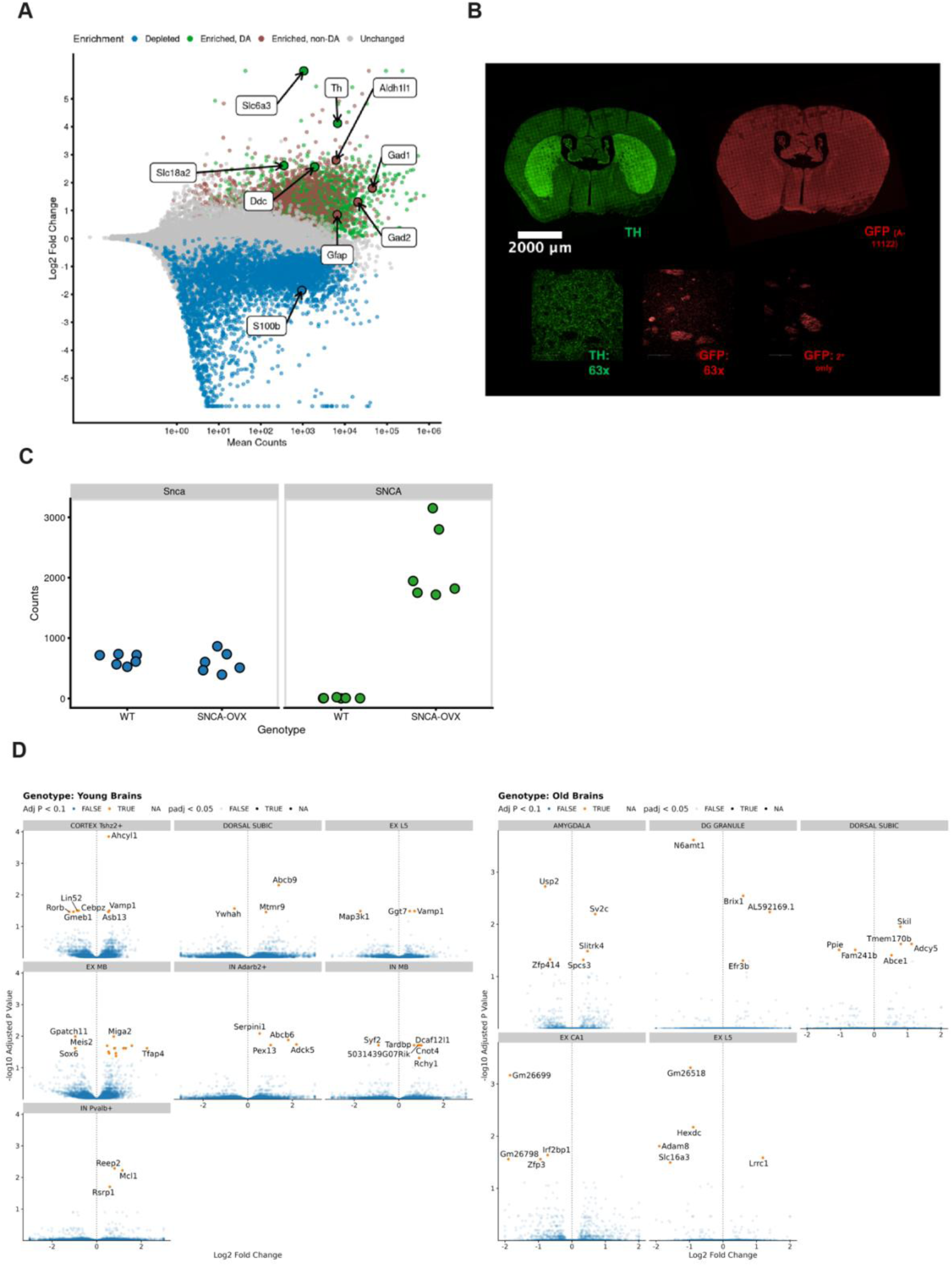
Investigating axonal translation in DA neurons and SNCA-OVX Characterization in TRAP and Stereo-seq samples. A) MA plot of enrichment/depletion in striatal TRAP samples. Markers of DA neurons (e.g. *Th, Slc6a3 (DAT), Slc18a2 (VMAT2)*) were enriched by TRAP, however markers of other cell types (e.g. *Gfap, Gad1*) were also enriched (*n* = 62 DAT-TRAP mRNA, n = 23 striatal homogenate mRNA). B) Immunohistochemical staining for GFP and TH in mouse striatum revealed the presence of faint puncta that colocalize with TH. C) Confirmation of *SNCA* overexpression specifically in *SNCA*-OVX samples (*n* = 12). *Snca* expression is unaffected by *SNCA* overexpression. Counts represent DESeq2-normalized read counts per sample. D) Volcano plots for all genes significantly differentially expressed due to *SNCA* overexpression across cell types of Stereo-seq samples.

